# Intracranial electrophysiology reveals distinct neural representations of nociception and salience in the human brain

**DOI:** 10.64898/2026.09.30.755799

**Authors:** Siddhartha Joshi, Mensure Polat, Sofia Pantis, Ritika Garg, Salaar Ahmed, Yuhao Huang, Paul D. McGeoch, Corey J. Keller, Vivek P. Buch, Sean Mackey, Josef Parvizi, Ashwin G. Ramayya

## Abstract

Neuroimaging studies have shown close overlap between neural representations of nociceptive stimuli and non-nociceptive, salient stimuli (such as low-probability sounds), leading to the proposal that nociception and salience share a common neural substrate in the human brain. Here we use high-resolution intracranial recordings from 13 neurosurgical patients (2,018 total electrodes) to show that neural representations of nociception and salience are anatomically and functionally distinct in the human brain. 77% of 107 electrodes that encoded nociceptive stimuli did not encode low-probability sounds (“nociceptive representations”), whereas 23% encoded both nociceptive and low-probability sounds (“salience representations”). Nociceptive representations were distinct from salience representations in terms of anatomical prevalence (posterior insula, inferior parietal, and perirolandic cortex vs. anterior insula, thalamus, and inferior frontal gyrus), timing (nociception preceded salience by 120 ms on a trial-by-trial basis), and function (intracranial electrical stimulation was more likely to cause sensory experience when applied near nociceptive representations, as compared to salience representations). These results provide strong evidence that nociception and salience have distinct neural substrates in the human brain, and that selective measurement of these distinct processes may help understand patient-to-patient variability in chronic pain and develop personalized brain stimulation treatments.

## Introduction

Understanding how the human brain processes nociceptive stimuli is important for improving diagnosis and treatment of chronic pain, which is a major problem in society (Lindsay et al., 2021). Previous studies using techniques such as scalp electroencephalography (EEG) or functional magnetic resonance imaging (fMRI) have identified neural representations of nociceptive stimuli in a wide range brain regions including somatosensory, cingulate, frontal and parietal cortices and the insula (Talbot et al., 1991; Ingvar, 1999; Peyron et al., 2000; Rainville, 2002; Tracey and Mantyh, 2007). However, the specificity of these neural representations for pain have been called into question because of the confound of salience (Legrain et al., 2011; Lee et al., 2020).

Salience is a property of a sensory stimulus which indicates how distinct it is from the environment and is thought to evoke several automatic neural processes including increased attention, preparatory motor activity, and sympathetic arousal. Because nociceptive stimuli are highly salient, their neural representations might reflect a general representation of salience that is not specific for nociception. Supporting this view, neural representations of nociceptive stimuli are closely overlapping with neural representations of non-nociceptive, salient stimuli, often studied using an auditory oddball paradigm which compares low-probability and standard tones (Mouraux et al., 2011; Horing et al., 2019; and Lee et al., 2020) and are largely unchanged in individuals with congenital insensitivity to pain (Salomons et al., 2016). These findings have led to the proposal that nociception and salience share a common neural substrate in the human brain, and that the brain processes nociceptive stimuli using the same neural populations that process salient stimuli across many sensory modalities (Legrain et al., 2011). However, more recent studies have shown that neural representations of nociception can be dissociated from salience within the posterior insula and parietal operculum using advanced statistical techniques, suggesting that neural representations of sensory pain and salience may be dissociated with higher resolution neural recordings (Horing et al., 2022).

Taken together, these observations have led to two competing hypotheses regarding human brain mechanisms for pain processing: first, that nociception and salience share a common neural substrate (Legrain et al., 2011), or second, that nociception and salience are processed by distinct neural populations that are activated in close spatial and temporal proximity, which leads to overlapping representations in functional neuroimaging studies. Here, we investigated these competing hypotheses using intracranial electroencephalography (iEEG) in 13 neurosurgical patients undergoing monitoring for medically refractory epilepsy as they performed two tasks: (1) A thermosensory nociceptive task (nociceptive vs. non-nociceptive thermal stimuli defined based on individual pain thresholds) and (2) a standard auditory oddball task (low– vs. high–probability tones). We were motivated by prior observations that direct measurements of neuronal activity with high spatiotemporal iEEG can distinguish between closely related neural representations that are spatially intermixed across the brain (Ramayya et al., 2015; 2025). We measured neural activity using high-frequency activity (HFA, 70–200 Hz power), which provides a reliable surrogate of local (within ∼3 mm) neural population spiking activity (Dubey and Ray, 2019; Manning et al., 2009; McCarty et al, 2022; Ray and Maunsell JH, 2011). We found that nociceptive and salience neural representations are largely distinct in the human brain. 77% of 107 electrodes that encoded thermosensory painful stimuli did not encode low-probability sounds (“nociceptive representations”), whereas 23% encoded both thermosensory nociceptive stimuli and low-probability sounds (“salience representations”). Nociceptive representations were distinct from salience representations in terms of anatomical prevalence, timing and function (as causally assessed by intracranial electrical stimulation, which was performed in a subset of patients).

## Methods

### Participants

In this study, we recruited 13 patients with medically refractory epilepsy who underwent surgical implantation of intracranial electrodes for seizure localization. Patients volunteered to perform cognitive testing as part of our research study and provided informed consent, while they were admitted to the hospital. These patients will hereby be referred to as “participants”. The study was approved by the Stanford University Institutional Review Board, and all procedures complied with relevant ethical regulations governing human subject’s research. Clinical considerations alone determined the number and anatomical placement of electrodes implanted, and their anatomical locations are shown in Figure 1A.

**Figure 1.**
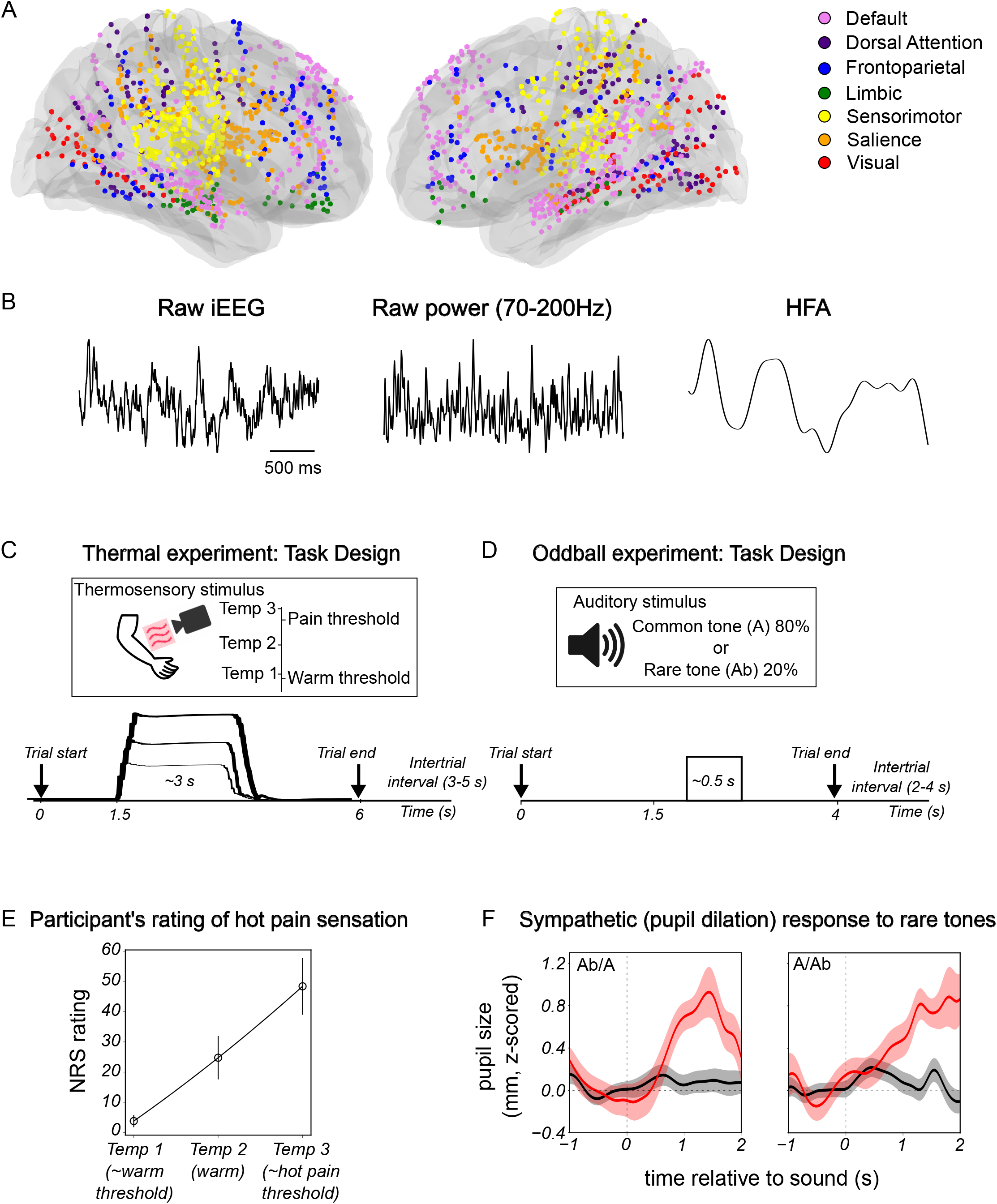
Task design. (A) Location of all 2,018 recording sites. (B) Intracranial EEG recording using depth electrodes: estimation of high frequency activity (HFA) as a surrogate for local neuronal spiking. (C) Thermal nociceptive task design. (D) Oddball task design. (E) Numerical ratings of pain. (F) Oddball experiment: example of pupil responses to tones.

### Thermal experiment

The thermal task (Figure 1C) was created and deployed using a TSA2/CHEPS system from Medoc, controlled via a laptop (Dell). Synchronizing timing signals (TTL pulses) were sent directly to the clinical EEG system while task and timing data were simultaneously saved on the laptop hard drive. The thermode was positioned on the participant’s arm, after which individual thresholds for warm detection, heat pain, cold detection, and cold pain were determined. Based on these thresholds, three stimulus intensities were selected for each participant: a temperature near the threshold for warm detection (low intensity; Temp 1), a non-nociceptive heat stimulus just below the heat pain threshold (medium intensity; Temp 2), and a nociceptive heat stimulus slightly above the heat pain threshold (high intensity; Temp 3; Figure 1C). Thermal stimuli were presented in randomized order using variable interstimulus intervals that depended on each participant’s response time, ensuring that subsequent trials began only after completion of the preceding response. Following each stimulus, participants rated perceived pain on a 0–100 numerical rating scale (NRS; Figure 1E). Representative recording locations and corresponding neural responses to thermal stimulation are shown in Figures 2A,B,G and H.

**Figure 2.**
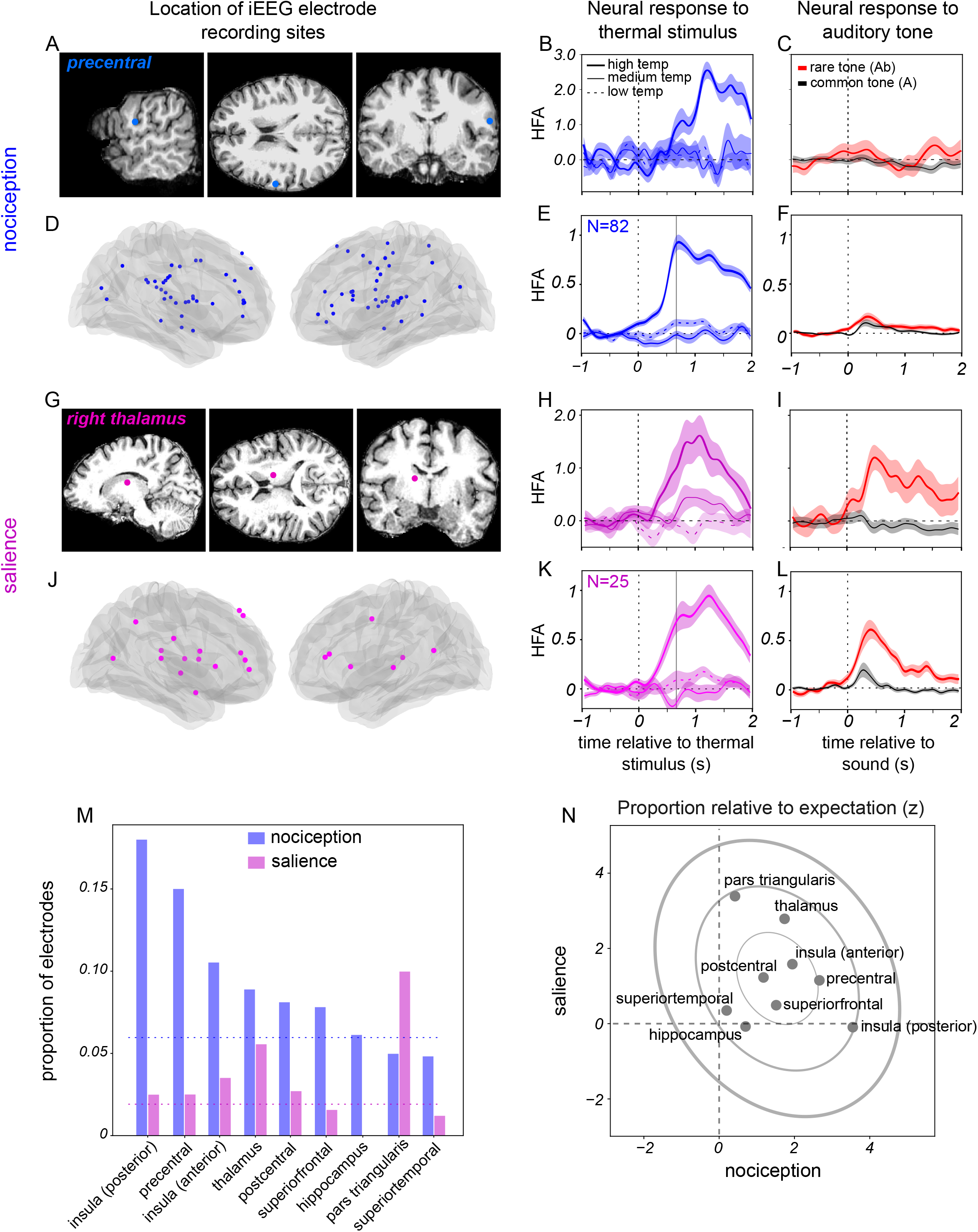
Most nociception encoding sites do not encode salience. (A) Location in the precentral gyrus of an electrode that was responsive to hot, painful stimulation only. HFA responses to (B) nociceptive stimuli and (C) auditory tones from this electrode. (D) Anatomical locations of all electrodes with nociceptive representation. Population averaged HFA responses to (E) nociceptive stimuli and (F) to tones for these electrodes. Salience responses: (G-I) Similar to A-C but now for a recording site in the thalamus that was responsive to both thermosensory nociceptive stimuli and low-probability sounds (salience representation). (J-L) similar to (D-F) but now for all sites responsive to both thermosensory painful stimuli and low-probability sounds. (M) Proportion of electrodes distributed among brain regions. (N) Relative frequency of sites identified as salience (ordinate) or nociceptive (abscissa).

### Auditory oddball experiment

The oddball task (Figure 1D) was created and deployed using Python and PsychoPy running on a laptop (Dell). In order to synchronize intracranial neural recordings with task events, timing signals were sent via a Labjack T4 device to the clinical EEG system. Each session started with verbal instructions given to the participant. Participants were asked to maintain their gaze on a fixation cross displayed at the center of a laptop screen, keep count of the number of oddball stimuli (low-probability A or A_b_ tones; 20% of all trials in a block) or the total number of tones presented, and report this number at the end of each block of trials. We consider the low-probability tones to be salient because of their statistical rarity, i.e. the tone was perceptually salient. These instructions were also displayed as text on the laptop screen prior to the appearance of the fixation cross. The stimulus timing and intertrial interval were both randomized to diminish stimulus timing predictability with interstimulus times varying between 2.5 and 4 sec. In each session, we ran two blocks of 100 trials each, switching the oddball tone in the second block (A_b_ to A, or A to A_b_). This switch was used to ensure that enhanced responses to the oddball tone were driven by its statistical rarity rather than by neural selectivity for a particular tone frequency. Consistent with previous work relating pupil size with stimulus salience and autonomic tone, we found that low-probability tones evoked larger pupil dilation compared with standard tones (Figure 1E; Murphy et al, 2011; Joshi et al., 2016). A site unresponsive to either common or low-probability tones is shown in Figure 2A with the response in Figure 2C, and a different, responsive site is shown in Figure 2G with the corresponding response in Figure 2I.

### Intracranial neural recordings

Intracranial EEG was recorded via depth electrodes (Ad-Tech or PMT Corporation) using the Nihon Kohden recording system with signals sampled at 1-2kHz. Signals from contacts on a single electrode were converted to a bipolar montage by taking the difference of signals on immediately adjacent contacts. The resulting bipolar signals were treated as new virtual electrodes (referred to as “electrode” hereafter) with location at the midpoint of each pair. We excluded electrodes with prominent line noise (60Hz), defined as greater spectral power in the 58-62Hz range than in the 18-22 Hz range, or electrodes that were disconnected (signal standard deviation=0).

### Intracranial electrical stimulation (iES)

Intracranial electrical stimulation was performed as part of routine clinical evaluation during presurgical epilepsy monitoring. The detailed protocol is available elsewhere (Parvizi et al., 2026). Briefly, bipolar stimulation was delivered between pairs of adjacent electrode contacts using a Nihon Kohden cortical stimulator (MS-120BK-EEG) with PE-210AK switchbox, interfaced with Nihon Kohden video-EEG monitoring equipment (WEE-1200, 1000 Hz sampling rate). Stimulation consisted of alternating square-wave pulses at 50 Hz frequency with 0.2 ms pulse width, delivered for 1-4 seconds per trial. Current amplitude was systematically increased but remained below the threshold for producing after-discharges, which was determined individually for each electrode pair. The patient was alerted by the physician prior to stimulation. Following each stimulation, the physician asked the patient what, if anything, they might have sensed. For example:

Physician: “How about now?”

Patient: “Ah there you go.”

Physician: “Yes, what happened?”

Patient: “My right hand started to feel heated and numb.”

### Anatomical localization of electrodes

Intracranial electrodes were identified manually on each post-operative CT scan, and contact coordinates were recorded based on the center of density of the radiodense contacts on thresholded images. Postoperative CT images were registered to each participant’s preoperative MRI, which was subsequently registered to the Montreal Neurological Institute (MNI) template to obtain electrode coordinates in both native and standardized anatomical space. We assigned each electrode to various canonical intrinsic brain networks (“7 network model”) using a volumetric atlas in MNI coordinates (Yeo et al., 2011). We refer to the “ventral attention” network as the “salience” network, but otherwise use terminology as reported in the original study.

### Extracting high-frequency activity (HFA)

We extracted three seconds of iEEG data around each thermal and auditory stimulus presentation, one second preceding the stimulus and two seconds following it. We defined this 3 second window as one trial for further analysis. We first used Welch’s method to estimate power spectral density between 0 and 500 Hz. We identified and rejected bad and disconnected electrodes as defined above (“Intracranial neural recordings”). We then extracted high-frequency spectral power for each trial using the continuous wavelet transform (PyWavelets: Lee et al., 2019; Figure 1B). We used a Morlet wavelet with bandwidth parameter 1.5 and center frequency 1.0 to optimize the tradeoff between time and frequency resolution, and logarithmically spaced frequencies between 70 and 200 Hz (Figure 1B). We then squared the wavelet convolution, log transformed the resulting power values, smoothed with a Gaussian filter (σ=30 ms) and then standardized each trial time-series by z-scoring using the mean and standard deviation from all trials in the block followed by subtracting the average activity just preceding each tone (baseline subtraction). Hereafter, we refer to this log-transformed, z-scored, baseline-corrected HFA as “HFA”.

### Analyzing responses to thermal and auditory stimuli

After examining individual trial HFA and per-electrode, trial averaged values, we defined response windows following every stimulus. For each trial, the HFA within this window was averaged to obtain the response to the tone or to the thermal stimulus. For the oddball experiment, for each electrode, we used a non-parametric test to determine whether the low probability tone reliably evoked larger responses than the standard tone (*H*_*0*_: response to low–probability tone ≤ response to standard tone, Mann-Whitney U-test, p<0.05). We used a similar approach to identify thermosensitive responses, except that HFA in the response window was compared with baseline activity preceding each high temperature stimulus. These electrodes were defined as “responsive”. The remaining electrodes were termed as “unresponsive”. To identify nociceptive representations among the thermosensitive group, we compared HFA responses evoked by stimuli just below pain threshold (medium temperature) with those evoked by stimuli at or above pain threshold (high temperature) stimuli (*H*_*0*_: response to high temperature ≤ response to medium temperature, Mann-Whitney U-test, p<0.05).

### Cross-correlation analysis

We performed a cross-correlation analysis to assess the temporal relation between the nociceptive and salience groups. For each trial, we averaged responses from all electrodes in each cluster to obtain a per-trial group-averaged response. Thus, for each participant, for each thermosensory stimulus, there was a single response for the nociceptive group and a single response for the salience group. We then calculated the trial cross-correlation between the two groups. We estimated noise correlations using time windows that excluded stimulus-evoked responses. For each trial, we subtracted the noise correlation from each trial’s signal correlation at each lag (“corrected” signal correlation). We then averaged the corrected signal correlation across all trials. 95% confidence intervals were computed from the aggregate distributions of noise-subtracted time series across each pair of electrodes. The same method was used to estimate the temporal relationship between thermal ramps (gradually increasing temperature; Figure Supplement 3A,B) and neural activity for the threshold experiment data.

### ROC analysis

For every electrode, we used the per-trial neural responses to medium (just below pain threshold) and high (hot, above pain threshold) temperature stimuli to estimate a receiver operating characteristic (ROC) curve (Green and Swets, 1965; Barlow, 1972; Britten et al, 1992). We calculated the area under each ROC curve (AUC) as a measure of neural discriminability between the medium and high temperatures.

### Intracranial Electrical Stimulation (iES) analysis

We determined the proportion of electrodes at which iES evoked a patient report of sensory experience. To increase our sample size for this analysis, we broadened our criteria used to select salience electrodes to include all electrodes at which we measured a selective response to low-probability sounds and an increase following the high temperature stimulus as compared to baseline (but did not require that the electrode show greater activity following high temperature than medium temperature stimuli).

### Code

We wrote custom code in Python for all data analysis. We used open-source scientific libraries and repositories including NumPy, SciPy, Scikit-learn, Pandas, MNE, and Nilearn. Code for the auditory oddball experiment was written in Python and PsychoPy. All code and data for this report are available at: https://github.com/RamayyaLab.

## Results

### Nociceptive and salience representations are anatomically distinct in the human brain

We measured high-frequency activity (HFA, 70-200 Hz power, henceforth “HFA”) as a surrogate of local population spiking activity from 2018 total electrodes across widespread brain regions (henceforth, “sites”; Figure 1). We identified 107 sites (9 participants) that showed increased HFA following painful stimuli. We subdivided these sites into two groups: “nociceptive representations,” if they were activated following painful stimuli, but not nonpainful salient stimuli (low-probability tones; 77%; 82/107), and “salience representations,” that were activated following both painful stimuli and low-probability tones (23%; 25/107). An example of a nociceptive representation is shown in Figure 2, demonstrating a robust response (Figure 2B) to thermosensory painful stimuli and no response (Figure 2C) to low-probability auditory stimuli. A different site showed salience representation, with clear responses to both painful stimuli (Figure 2H) and low-probability auditory stimuli (Figure 2I).

Both nociceptive and salience representations were widely distributed throughout the brain (Figure 2M; Figure Supplement 1, Table 1). Nociceptive representations were prominent in many regions identified in prior neuroimaging studies including insula, pre- and post-central gyri, but also observed in regions that are not typically identified in human neuroimaging studies, such as the pars triangularis region of the inferior frontal gyrus. We observed salience representations in a smaller set of regions, including anterior insula and thalamus. When considering distributions across intrinsic brain networks, nociceptive representations were distributed mainly within the sensorimotor network while salience representations were predominantly observed within the salience (“ventral attention”) network (Figure Supplement 1 ).

**Table 1.**
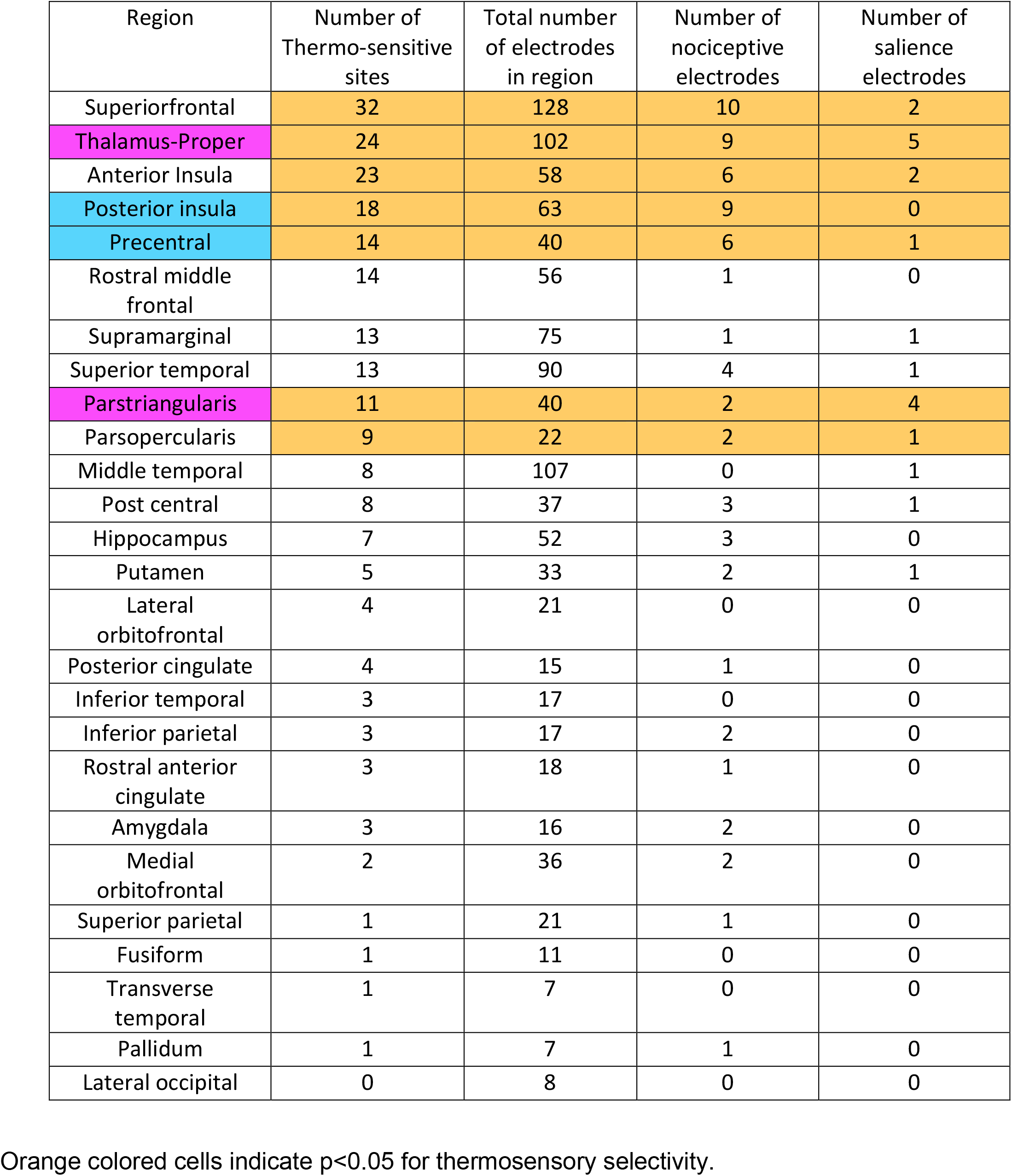
Anatomical distribution of recording sites and prevalence of nociceptive and salience electrodes. Orange colored cells indicate p<0.05 for thermosensory selectivity.

### Nociceptive representations are functionally connected to and precede salience representations in the human brain

We estimated the relative timing between nociceptive and salience representations using cross correlation analysis on the responses to thermal ramp and hold stimuli (Figure 1C) and found that nociceptive representations were correlated with salience representations and preceded them by 120ms (Figure 3A). Consistent with these faster sensory pain responses, we found that responses to slow thermal ramps (used in the thermal threshold experiment) were more reliably correlated (H_0_: correlation between temperature and HFA is>0, t-test p<0.05) with increasing temperature for the pain group (Figure Supplement 2 A,C) than for the salience group (Figure Supplement 2 B,D).

**Figure 3.**
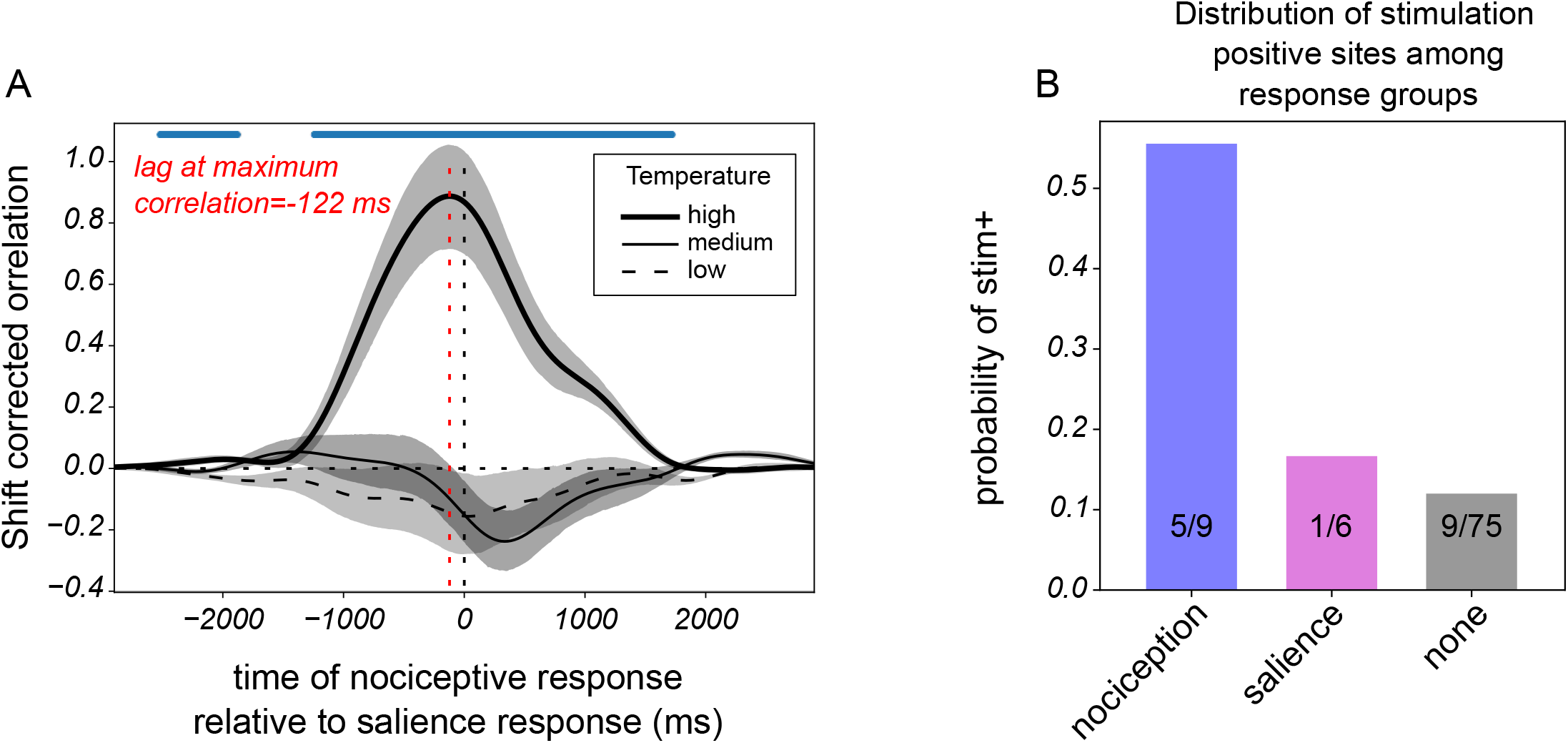
Nociception related neural activity precedes salience activity and iES evokes stronger thermosensation in the nociceptive group than the salience group. Relative timing of neural responses following thermal stimuli from nociceptive and salience electrodes. (A) Cross correlation between nociceptive and salience responses to thermosensory ramp-and-hold stimuli at high (heavy line) medium (light line) and low (dashed line) temperatures. Red vertical dashed line indicates time of peak correlation for the high temperature condition. Shaded regions show estimated 95% confidence intervals. (B) Relating subjective responses during iES with neural responses evoked by thermosensory painful stimulation: distribution of stimulation positive sites among the response groups. The number of stimulated sites within each group is noted within each bar.

### Nociceptive representations differ from salience representations in causal effects on sensory experience

We found that intracranial electrical stimulation near electrodes that showed nociceptive representations were more likely to cause subjective reports of sensory experience as compared to electrodes that showed salience representations (Figure 3B). In a subset of sites (N=132), we performed intracranial electrical stimulation (iES) and assessed subjective reports of sensory experience. iES is thought to modulate local neural activity and can reveal causal relations between local neural activity and experience (Duong et al., 2023). We classified the iES sites as those that were “positive” (i.e., evoked a report of a sensory experience, including painful and thermosensory experiences) and those that were “negative” (that did not evoke such an experience). In all, we observed 21/132 positive sites and found that electrodes that showed nociceptive representations were more likely to be classified as positive sites than electrodes that showed salience representations (Figure 3B; 56%, 5/9, vs. 17%, 1/6, respectively, Binomial test, z=3.13; *p* =0.009). Of note, we broadened our definition of salience representations for this analysis to increase our sample size for this subgroup analysis (see Materials and Methods). Among all positive iES sites, 6/21 were located outside of the posterior insula, a region with a strong representation of thermal pain (Figure 2M,N; 3 in posterior cingulate, 2 in superior frontal and one in postcentral cortex).

### Both nociceptive and salience representations correlate with trial-to-trial variability in pain reports and showed similar strength of pain encoding

Both nociceptive and salience representations covaried with subjective reports of pain. Responses to medium and high temperature thermosensory stimuli from a site with nociceptive representation are shown in Figure 4A and B, respectively. We estimated slopes from linear regression on the data (lines in Figure 4A and B). Mean slopes and 95% confidence intervals for the two groups are shown in Figure 4C, demonstrating a positive relationship between neural representation and trial-to-trial variability in pain reports for both groups.

**Figure 4.**
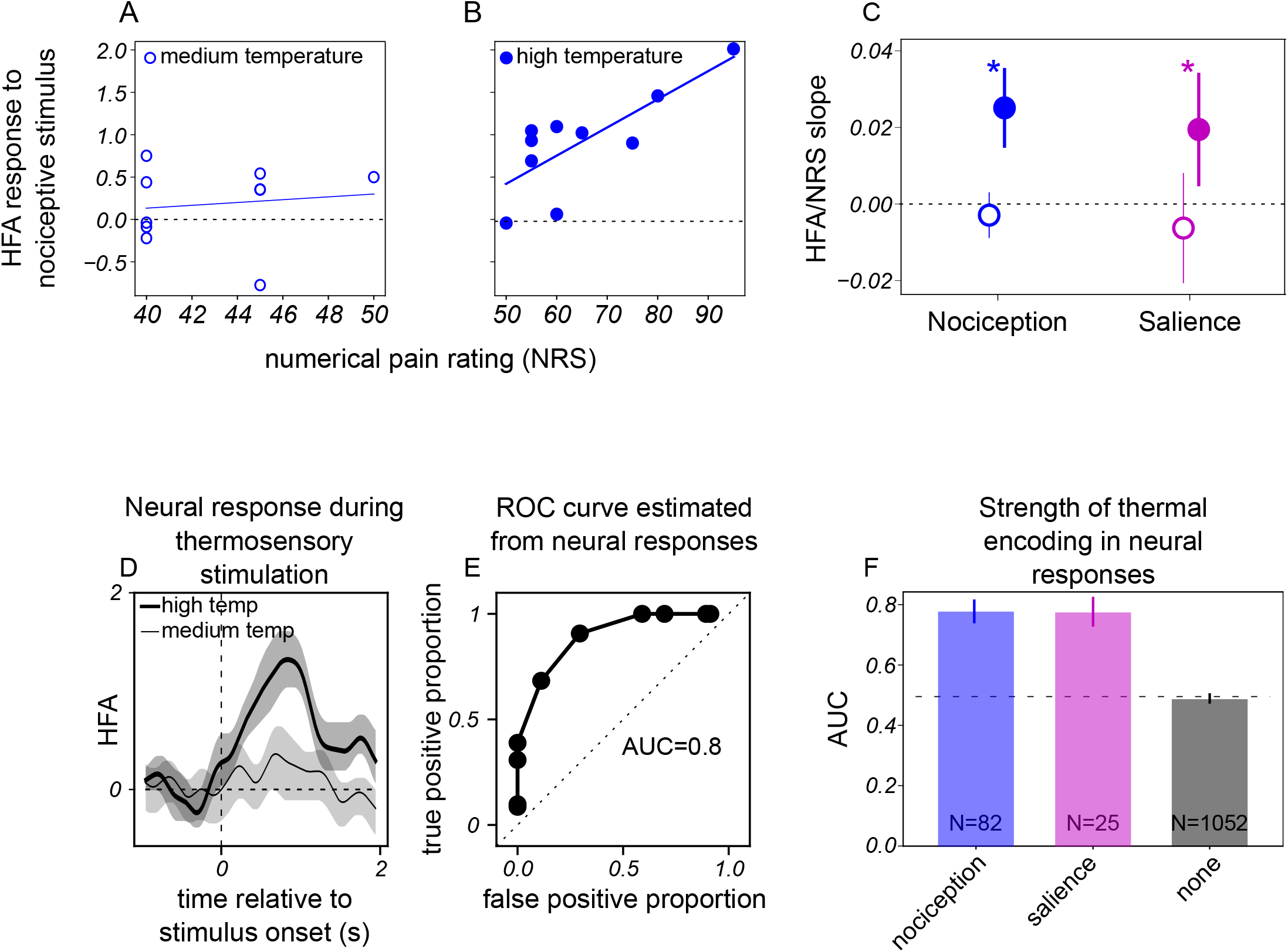
Neural responses corresponding to both nociceptive and salience activations correlate with pain experience and have similar encoding for thermal stimuli. Neural responses corresponding to both nociceptive and salience representations correlate with pain experience and have similar encoding for thermal stimuli. HFA responses versus pain ratings for (A) medium and (B) high temperature thermosensory stimuli recorded at a postcentral electrode. Lines in both panels are linear fits to the data points. (C) Mean and 95% confidence intervals of the slopes for all data from each group. Asterisks above markers indicate p<0.05 for *H*_*0*_: mean of medium temperature slopes is same as mean of high temperature slope (*t*-test). (D) HFA responses recorded at the same example electrode (shown in A and B) to thermosensory stimuli. (E) Discriminating between nociceptive and non-nociceptive thermosensory stimuli using neural responses: ROC curve estimated from neural responses shown in (D). (F) Mean and 95% confidence intervals for the area under the curve (AUC) estimated from neural responses.

In terms of encoding strength, we found that nociceptive and salience representations discriminated between nociceptive and non-nociceptive thermal stimuli with similar strength. We used HFA responses to thermosensory stimuli to estimate the strength of nociceptive encoding. An example site with a clear response to thermosensory painful stimuli is shown in Figure 4D, with a receiver operating characteristic (ROC) curve estimated from these responses shown in Figure 4E. For this example, the area under the ROC curve (AUC) was 0.8, indicating that the neural activity at this location could reliably distinguish between hot painful and warm stimuli. For each response group described above (also see Figure 2), the mean area under the ROC curve (AUC) is shown in Figure 4F. Neural activity at sites classified as nociceptive and salience showed AUC values that indicate discriminability between hot painful and warm stimuli (Figure 4F, blue and magenta bars). The discriminability of these two groups did not differ reliably (*H*_*0*_: nociceptive AUC values different from salience AUC values, Mann-Whitney U test, p=0.3). On the other hand, sites that were unresponsive to either stimulus showed poor discriminability for thermal painful stimuli (Figure 3C, black bars) and were reliably poorer at such when compared with the nociceptive and salience groups (*H*_*0*_: nociceptive or multimodal salience AUC values different from rest of sites, Mann-Whitney U test, p<0.005).

## Discussion

We found that the human brain contains a distributed neural representation of nociception that is distinct from salience. Approximately 77% of neural populations that encoded nociception did not encode low-probability sounds, which are nonpainful, salient stimuli (nociceptive representations), whereas ∼23% encoded both painful stimuli and low-probability sounds (salience representations), indicating a neural representation of general salience which is not specific to pain. Nociceptive and salience representations were distinct by three measures: first, nociceptive representations showed distinct anatomical distribution than salience representations (posterior insula, inferior parietal, and perirolandic cortex vs. anterior insula, thalamus, and inferior frontal gyrus). Second, nociceptive representations occurred 120 ms earlier than salience on a trial-by-trial basis per a cross-correlation analysis. Third, intracranial electrical stimulation was more likely to evoke subjective reports of sensory experience when applied near nociceptive representations as compared to salience representations, providing causal evidence for a functional dissociation between these neural populations. Taken together, these results challenge the view that nociception and salience share a common neural substrate in the human brain but instead support competing view that human brain contains distinct neural representations of nociception and salience, which are regionally overlapping, but functionally dissociable. Our finding of a sequential representation of nociceptive and salience neural populations is broadly consistent with contemporary views of how the human brain generates sensory experience, whereby unimodal sensory neural representations transform into integrative representations in higher order brain regions (e.g., nociception in posterior insula to salience in anterior insula; Hua, 2005; Craig, 2009; Rao and Ballard, 1999; Friston, 2009).

How might we reconcile our findings with prior non-invasive neuroimaging studies which show overlapping representations between nociception and salience? One possibility is that our intracranial high frequency activity measurements, which provides an index of local spiking activity with high spatiotemporal resolution (∼50 ms and ∼ 3mm; Manning et al., 2009; Ray and Maunsell, 2011; Dubey and Ray 2019), provides a more detailed view of local neural representations in the human brain. Prior studies have shown that these measurements can resolve functionally distinct neural populations which are spatially intermixed within a brain region in a way that is challenging for aggregate region-level neural measurements (Ramayya et al., 2015; 2025). Non-invasive neuroimaging in past studies could, in part, limit their ability to resolve small populations of neurons with a given region. Specifically, scalp EEG measurements have limited spatial resolution that makes it difficult to disentangle neural populations for nociception and salience (Mouraux et al., 2011, Salomons et al., 2016; although, see Horing et al., 2019, 2022). As we demonstrated using iES in the insula, even within a given brain region that is co-activated by nociceptive and non-nociceptive salient stimuli, there might be spatially segregated heterogenous neural populations specific either for nociception or for multimodal salience, that very likely are aggregated together by scalp EEG and fMRI studies. The fMRI technique has additional limitations including that: (i) it is an indirect measurement of neural activity, (ii) it likely reflects aggregated responses over relatively large spatial scales and (iii) might sometimes even be dissociated from local neuronal activity (Logothetis, 2003; Ekstrom, 2010; Howarth et al., 2021).

Our results provide novel insights into how the human brain processes nociception in relation to salience. Whereas previous studies have used intracranial recordings to investigate nociceptive processing in the human brain, they have typically focused on specific brain regions and/or used analyzed evoked potentials or low frequency activity, which are non-specific indicators of local spiking activity (Liberati et al., 2016, 2017, 2019, 2020; Soyman et al., 2022; Shivralkar et al, 2023), or did not did not investigate the relationship between nociception and salience (Bastuji et al., 2016; Caston et al., 2023).

Our study has several limitations. First, it is possible that some of these results are particular to our patient population. In general, epilepsy patients can show additional forms of inter-individual variability in their brain networks related to their pathology that might also manifest in certain categorical behavioral differences compared to healthy controls. However, we sought to mitigate such concerns by focusing on neural signals associated with specific, highly controlled stimuli. Second, given our vulnerable patient population and the clinical setting, we only provided mildly painful stimuli for ethical reasons, which limited the range of pain ratings that we observed during the experiment. Third, because electrode implantation was purely driven by clinical indications, we had sparse anatomical sampling throughout the brain.

We found that the human brain sequentially encodes nociception and salience via anatomically and functionally distinct neural populations. These results argue against the prevailing view that nociception is primarily processed through a general salience circuit and instead suggest that nociception and salience are distinct and identifiable neural representations in the human brain. Future studies should investigate how these distinct representations are integrated together generate pain, and whether selectively targeting these distinct neural populations can fuel advances in diagnostic and treatment approaches for patients with chronic pain.

## Acknowledgements

We thank Jiaqi Wu and Creence Lin for their expert help in developing experiment task code, and Zoe Lusk for expert help in collecting and curating iEEG data.

---

**Figure Supplement 1.**
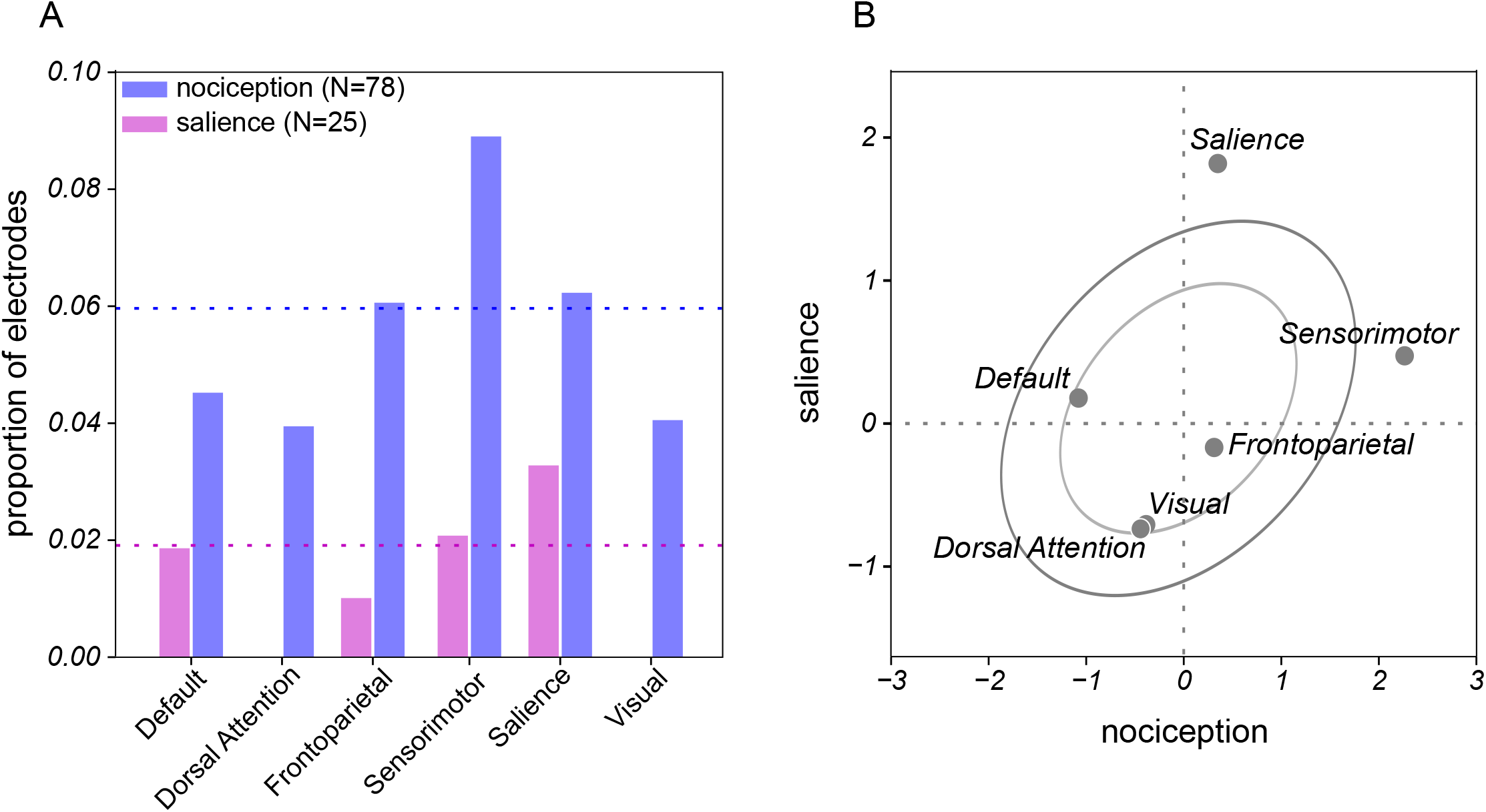
Brain networks for nociception and salience. (A) Distribution of electrodes among standard brain networks. (B) Relative frequency of salience (ordinate) or nociceptive representations (abscissa; z-scores on both axes).

**Figure Supplement 2.**
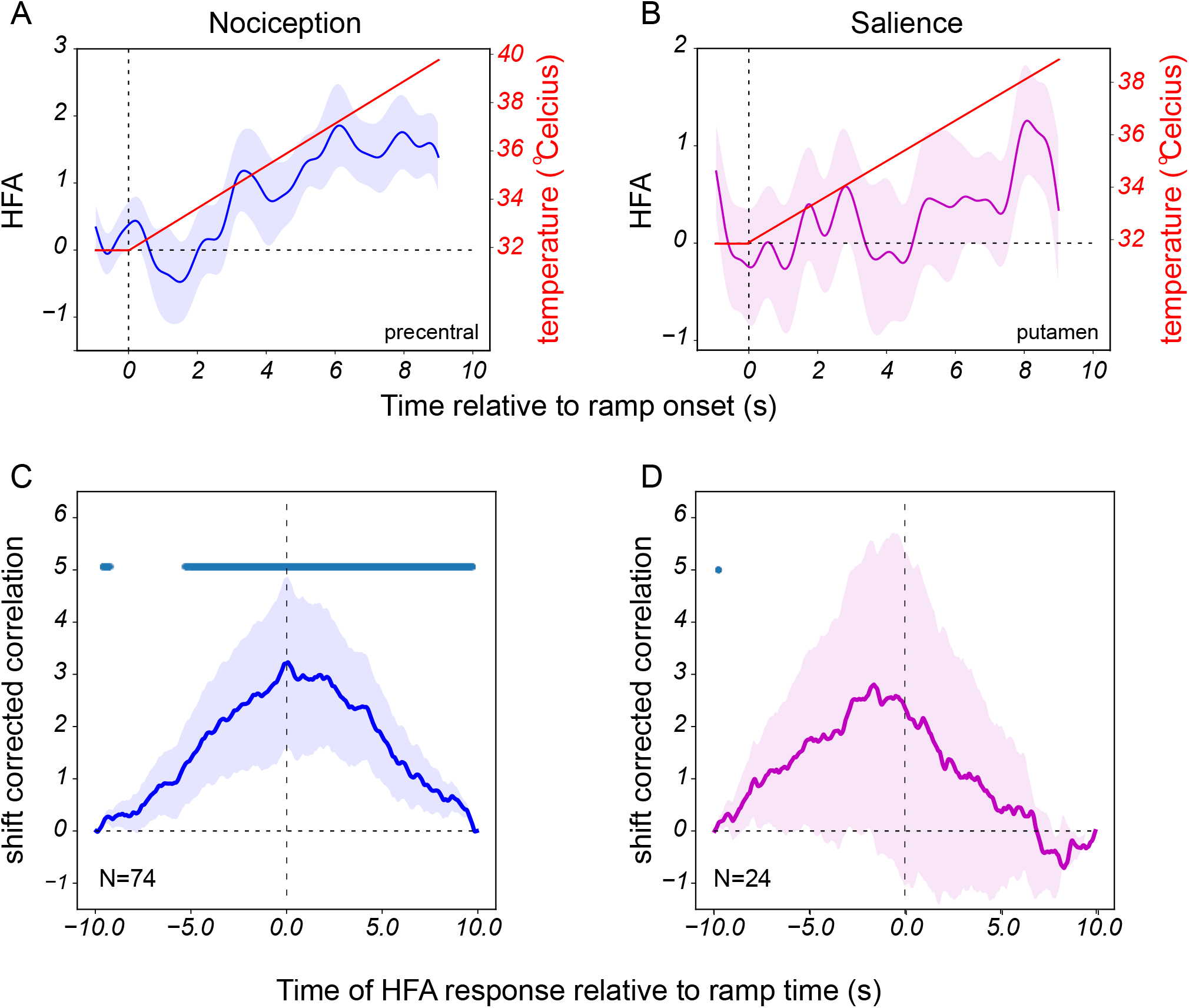
Correlation between thermal ramp stimulus and neural response for the two response groups. Example neural responses to thermosensory ramps (shown in red) from the (A) nociceptive and (B) salience groups. The HFA is aligned to time of onset of thermosensory ramp . (C) and (D) Correlation between temperature and HFA response for the two groups. Markers above correlogram indicate p<0.05 for *H*_*0*_: correlation>0, t-test.

## Notes

### Competing Interest Statement

The authors have declared no competing interest.

### Funding

- NIH: R21NS145164, R21NS140918

## References

1. Barlow HB. Single units and sensation: a neuron doctrine for perceptual psychology? Perception. 1972;1(4):371–394.

2. Bastuji H, Frot M, Perchet C, Magnin M, Garcia-Larrea L. (2016). Pain networks from the inside: Spatiotemporal analysis of brain responses leading from nociception to conscious perception. Hum Brain Mapp. 37:4301–4315.

3. Britten KH, Shadlen MN, Newsome WT, Movshon JA. (1992). The analysis of visual motion: a comparison of neuronal and psychophysical performance. J Neurosci. 12:4745 –4765.

4. Caston RM, Smith EH, Davis TS, Singh H, Rahimpour S, Rolston JD. (2023). Characterization of spatiotemporal dynamics of binary and graded tonic pain in humans using intracranial recordings. PLoS One. 18(10):e0292808.

5. Craig AD. (2009). How do you feel--now? The anterior insula and human awareness. Nat Rev Neurosci. 10:59–70.

6. Dubey, A. & Ray, S. Cortical electrocorticogram (Ecog) is a local signal. Journal of Neuroscience 39, 4299–4311 (2019).

7. Duong A, Quabs J, Kucyi A, Lusk Z, Buch V, Caspers S, Parvizi J. (2023). Subjective states induced by intracranial electrical stimulation matches the cytoarchitectonic organization of the human insula. Brain Stimul. 16:1653–1665.

8. Ekstrom A. (2010). How and when the fMRI BOLD signal relates to underlying neural activity: the danger in dissociation. Brain Res Rev. 62:233–244.

9. Friston, K. The free-energy principle: a unified brain theory?. Nat Rev Neurosci 11, 127– 138 (2010). 10.1038/nrn2787

10. Green DM, Swets JA. (1966). Signal detection theory and psycho-physics. New York: Wiley.

11. Horing, B., Sprenger, C., & Büchel, C. (2019). The parietal operculum preferentially encodes heat pain and not salience. PLoS biology, 17(8), e3000205.

12. Horing B, Büchel C. (2022). The human insula processes both modality-independent and pain-selective learning signals. PLoS Biol. 20(5):e3001540.

13. Howarth C, Mishra A, Hall CN. (2021). More than just summed neuronal activity: how multiple cell types shape the BOLD response. Philos Trans R Soc Lond B Biol Sci. 376(1815):20190630.

14. Hua le H, Strigo IA, Baxter LC, Johnson SC, Craig AD. (2005). Anteroposterior somatotopy of innocuous cooling activation focus in human dorsal posterior insular cortex. Am J Physiol Regul Integr Comp Physiol. 289(2):R319–R325.

15. Ingvar, M. (1999). Pain and functional imaging. Philos. Trans. R. Soc. Lond. B: Biol. Sci. 354, 1347–1358.

16. Joshi S, Li Y, Kalwani RM, Gold JI. (2016). Relationships between Pupil Diameter and Neuronal Activity in the Locus Coeruleus, Colliculi, and Cingulate Cortex. Neuron. 89:221–234.

17. Lee, I. S., Necka, E. A., & Atlas, L. Y. (2020). Distinguishing pain from nociception, salience, and arousal: how autonomic nervous system activity can improve neuroimaging tests of specificity. Neuroimage, 204, 116254.

18. Legrain, V., Iannetti, G. D., Plaghki, L., & Mouraux, A. (2011). The pain matrix reloaded: a salience detection system for the body. Progress in neurobiology, 93(1), 111 –124.

19. Lindsay, N. M., Chen, C., Gilam, G., Mackey, S. & Scherrer, G. (2021). Brain circuits for pain and its treatment. Science Translational Medicine vol. 13 Preprint at 10.1126/scitranslmed.abj7360

20. Logothetis NK. (2003). The underpinnings of the BOLD functional magnetic resonance imaging signal. J Neurosci. 23:3963–3971.

21. Manning JR, Jacobs J, Fried I, Kahana MJ. (2009). Broadband shifts in local field potential power spectra are correlated with single-neuron spiking in humans. J. Neurosci. 29, 13613–13620.

22. McCarty MJ, Woolnough O, Mosher JC, Seymour J, Tandon N. (2022). The Listening Zone of Human Electrocorticographic Field Potential Recordings. eNeuro. 9(2):ENEURO.0492 -21.2022.

23. Mouraux, A., & Iannetti, G. D. (2009). Nociceptive laser-evoked brain potentials do not reflect nociceptive-specific neural activity. Journal of neurophysiology, 101(6), 3258 –3269.

24. Mouraux, A., Diukova, A., Lee, M. C., Wise, R. G., & Iannetti, G. D. (2011). A multisensory investigation of the functional significance of the “pain matrix”. Neuroimage, 54(3), 2237–2249.

25. Murphy PR, Robertson IH, Balsters JH, O’connell RG. (2011). Pupillometry and P3 index the locus coeruleus-noradrenergic arousal function in humans. Psychophysiology. 48:1532–1543.

26. Parvizi J, Julian Quabs J, Pantis S, Chen G, Huang W, Ma E, Del Vecchio M, Lyu D, Wang C, Avanzini P, Buch V, Ramayya A, Caspers S, Vogel H. (2026). Functional Architecture of the Human Insula Revealed by Causal Intracranial Mapping. 10.21203/rs.3.rs-8912902/v1

27. Peyron, R., Laurent, B., Garcia-Larrea, L. (2000). Functional imaging of brain responses to pain. A review and meta-analysis Neurophysiol. Clin. 30, 263–288.

28. Rainville, P. (2002). Brain mechanisms of pain affect and pain modulation. Curr. Opin. Neurobiol. 12, 195–204.

29. Ramayya AG, Buch V, Richardson A, Lucas T, Gold JI. (2025). Human response times are governed by dual anticipatory processes with distinct neural signatures. Commun Biol. 8(1):124.

30. Ramayya AG, Pedisich I, Kahana MJ. (2015). Expectation modulates neural representations of valence throughout the human brain. Neuroimage. 115:214 –23.

31. Rao, R., Ballard, D. Predictive coding in the visual cortex: a functional interpretation of some extra-classical receptive-field effects. Nat Neurosci 2, 79–87 (1999). 10.1038/4580

32. Ray S, Maunsell JH. (2011). Different origins of gamma rhythm and high-gamma activity in macaque visual cortex. PLoS Biol. 9, e1000610.

33. Salomons TV, Iannetti GD, Liang M, Wood JN. (2016). The “Pain Matrix” in Pain-Free Individuals. JAMA Neurol. Jun 1;73(6):755–6. doi: 10.1001/jamaneurol.2016.0653.

34. Shirvalkar P, Prosky J, Chin G, Ahmadipour P, Sani OG, Desai M, Schmitgen A, Dawes H, Shanechi MM, Starr PA, Chang EF. (2023). First-in-human prediction of chronic pain state using intracranial neural biomarkers. Nat Neurosci. 26:1090–1099.

35. Soyman E, Bruls R, Ioumpa K, Müller-Pinzler L, Gallo S, Qin C, van Straaten ECW, Self MW, Peters JC, Possel JK, Onuki Y, Baayen JC, Idema S, Keysers C, Gazzola V. (2022). Intracranial human recordings reveal association between neural activity and perceived intensity for the pain of others in the insula. Elife. 11:e75197.

36. Talbot JD, Marrett S, Evans AC, Meyer E, Bushnell MC, Duncan GH. (1991). Multiple representations of pain in human cerebral cortex. Science. 251:1355–1358.

37. Tracey, I., Mantyh, P.W. (2007). The cerebral signature for pain perception and its modulation. Neuron 55, 377–391.

